# Rapid Characterization of Human Serum Albumin Binding for Per- And Polyfluoroalkyl Substances Using Differential Scanning Fluorimetry

**DOI:** 10.1101/2021.06.13.448257

**Authors:** Thomas W. Jackson, Chris M. Scheibly, M. E. Polera, Scott M. Belcher

## Abstract

Per- and polyfluoroalkyl substances (PFAS) are a diverse class of synthetic chemicals that accumulate in the environment. Many proteins, including the primary human serum transport protein albumin (HSA), bind PFAS. The predictive power of physiologically based pharmacokinetic modeling approaches are currently limited by a lack of experimental data defining albumin binding properties for most PFAS. A novel thermal denaturation assay was optimized to evaluate changes in thermal stability of HSA in the presence of increasing concentrations of known ligands and a structurally diverse set of PFAS. Assay performance was initially evaluated for fatty acids and HSA binding drugs ibuprofen and warfarin. Concentration response relationships were determined and dissociation constants (K_d_) for each compound were calculated using regression analysis of the dose-dependent changes in HSA melting temperature. Estimated K_d_ values for HSA binding of octanoic acid, decanoic acid, hexadecenoic acid, ibuprofen and warfarin agreed with established values. The binding affinities for 24 PFAS that included perfluoroalkyl carboxylic acids (C4-C12), perfluoroalkyl sulfonic acids (C4-C8), mono- and polyether perfluoroalkyl ether acids, and polyfluoroalkyl fluorotelomer substances were determined. These results demonstrate the utility of this differential scanning fluorimetry assay as a rapid high through-put approach for determining the relative protein binding properties and identification of chemical structures involved in binding for large numbers of structurally diverse PFAS.

## 1. Introduction

Per- and polyfluoroalkyl substances (PFAS) are a large class of persistent synthetic chemicals used in a wide-variety of industrial and consumer applications.^1–3^ The perfluorinated aliphatic backbones of PFAS are hydrophobic, chemically inert, and thermally stable; consequently, they are persistent and accumulate in the environment and in biota.^4^ The most recent comprehensive analysis by the Organization of Economic Cooperation and Development identified > 4,730 PFAS-related CAS registry numbers, including 947 compounds that were registered in the EPA Toxic Substances Control Act (TSCA) chemical inventory.^5^

Production and use of long-chain perfluoroalkyl acids (PFAA; e.g, perfluoroalkycarboxylic acids (PFCA) with ≥ seven fluorinated carbons and perfluoroalykylsulfonic acids (PFSA) with ≥ six fluorinated carbons), began in the 1950s and continued in the United States until 2002, when manufacturers began to phase out long-chain PFAA due to their persistence and toxicity. As a response to the phaseout, short-chain PFAS are increasingly used as replacement chemistries in many applications and processes.^6^ Common examples of these replacement chemistries include PFCA and PFSA with shorter fluoroalkyl chains [e.g. perfluorobutanecarboxylic acid (PFBA) and perfluorobutanesulfonic acid (PFBS)], per- and polyfluoroalkyl ether acids (PFEA) that contain one or more ether moieties [e.g. hexafluoropropylene oxide dimer acid (HFPO-DA)], and fluorotelomer acids and alcohols with perfluoroalkyl length ≤ six.^1,7,8^ Since their introduction, shorter chain replacement PFAS are now detected ubiquitously in the environment and are accumulating in people and other organisms across the world.^9–11^

The physiochemical properties, exposure, and toxicity of perfluorooctanoic acid (PFOA) and perfluorooctanesulfonic acid (PFOS) are most well characterized. By contrast, there are only limited data available for the majority of known PFAS, including most of the replacement PFAS currently in use. The 1000’s of PFAS for which there is a paucity of available data necessitates the use of high throughput and predictive computational strategies to characterize the physiochemical properties, bioactivity, and potential toxicity across different classes of PFAS. Recently, physiologically-based pharmacokinetic and molecular dynamics modeling, quantitative structure-activity relationship, and machine learning approaches have been developed to predict protein binding affinity for PFAS.^12,13^ The predictive capabilities of these approaches are currently limited by a lack of data defining fundamental physio-chemical and toxicokinetic properties for most PFAS.

Albumin, the primary transport protein for PFOS, PFOA, perfluorononanoic acid (PFNA), perfluorohexanesulfonic acid (PFHxS), and perfluorodecanoic acid (PFDA), contains multiple non-specific binding sites that selectively bind fatty acids, hormones, drugs, and some xenobiotics including PFAS.^14^ However, experimentally determined binding affinities of most PFAS at albumin are unavailable. Current approaches for determining protein binding affinities, including titration chemistry or surface plasmon resonance, are too resource intensive and time-consuming to individually determine albumin affinity for each of the thousands of different PFAS.

Differential scanning fluorimetry (DSF) is a rapid high throughput method for measuring ligand binding interactions that is most often used to assess protein stability under various conditions.^15–17^ The DSF assay employs an environmentally sensitive fluorophore that is quenched while free in solution. Binding of the dye to hydrophobic sites accessible as the protein unfolds as temperature rises causes unquenching and fluorescence proportional to the amount of bound dye.^18,19^ Protein binding of ligand causes a concentration- and affinity-dependent stabilization of the folded protein structure observed as an increase in the melting temperature (T_m_).^16,20,21^ Relative binding affinity of the stabilizing ligand can be calculated from the dose-response relationship for the change in the T_m_.^17^

The goals of this study were to develop and optimize a high throughput DSF assay to rapidly characterize the relative HSA binding affinity of a variety of different PFAS. An initial set of control compounds, including fatty acids and albumin-binding drugs ibuprofen and warfarin, which bind HSA at different binding sites, were used to demonstrate feasibility and evaluate whether binding affinities estimated from DSF were comparable to known values estimated by other methods. Following optimization of DSF for PFOA and PFOS, binding affinity at HSA was determined for a structurally diverse set of PFAS that included nine perfluoroalkyl carboxylic acids of increasing chain length (C4-C12), three perfluoroalkyl sulfonic acids, four ether containing PFAS and eight fluorotelomer substances. The results from these analyses reveal that DSF approaches can be used to define protein-binding affinities rapidly and accurately for large numbers of chemically distinct PFAS, and this approach is able to discriminate between structurally similar PFAS. These results provide essential experimental data to better understand this diverse group of environmental contaminants.

## 2. Materials and Methods

### 2.1 Chemicals and reagents

Reagents and solvents used were the highest purity available. All aqueous buffers and solutions were prepared in sterile Milli-Q A10 water (18Ω; 3 ppb total oxidizable organics). GloMelt (*λ*_Ex_ = 468 nm *λ*_Em_ = 507 nm) and carboxyrhodamine (ROX; *λ*_Ex_ = 588 nm *λ*_Em_ = 608 nm) dyes were purchased from Biotium (Fremont, CA). The PFAS analyzed are shown in Figure 1. Octanoic acid (CAS 124-07-2, purity ≥ 98%), Perfluorobutanoic acid (PFBA, CAS 375-22-4, purity ≥ 99%), perfluoropentanoic acid (PFPeA, CAS 2706-90-3, purity ≥ 97%), perfluoroheptanoic acid (PFHpA, (CAS 375-85-9, purity ≥ 98%), PFOA (CAS 335-67-1 purity ≥ 95%), perfluorodecanoic acid (PFDA, CAS 335-76-2, purity ≥ 97%), perfluorododecanoic acid (PFDoA, CAS 307-55-1, purity ≥ 96%), perfluorotetradecanoic acid (PFTDA, CAS 376-06-7, purity ≥ 96%), and HFPO-DA (CAS 13252-13-6, purity ≥ 97%) were from Alfa Aesar (Havermill, MA). Perfluorohexanoic acid (PFHxA, CAS 307-24-4, purity ≥ 98%), perfluorononanoic acid (PFNA, CAS 375-95-1, purity ≥ 95%), Perfluorobutanesulfonic acid (PFBS, CAS 375-73-5, purity ≥ 98%), Warfarin (CAS 81-81-2, purity ≥ 98%), and 1H, 1H, 2H, 2H-Perfluorohexane-1-ol (4:2-FTOH, CAS 2043-47-2, purity ≥ 97%) were from TCI America (Portland, OR). Perfluoroundecanoic acid (PFunDA, CAS 2058-94-8, purity ≥ 96%) was from Oakwood Chemical (Estill, SC), perfluorohexanesulfonic acid (PFHxS, CAS 3871-99-6, purity ≥ 98%) was from J&K Scientific (Beijing, China), and PFOS (CAS 2795-39-3, purity ≥ 98%) and Perfluoro-3,6,9-trioxadecanoic acid (PFO3DoDA, CAS 151772-59-7, purity 98%) were from Matrix Scientific (Columbia, SC). Nafion byproduct 2 (CAS 749836-20-2, purity ≥ 95%), 1,1,1,2,2,3,3-Heptafluoro-3-(1,2,2,2-tetrafluoroethoxy)propane (E1, CAS 3331-15-2, purity ≥ 97%), 1H, 1H, 2H, 2H-Perfluorooctanol (6:2-FTOH, CAS 647-42-7, purity ≥ 97%), 2H,2H,3H,3H-Perfluorohexanoic acid (3:3-FTCA, CAS 356-02-5, purity ≥ 97%), 2H,2H,3H,3H-Perfluorooctanoic acid (5:3-FTCA, CAS 914637-49-3, purity ≥ 97%), 2,H,2H,3H,3H-Perfluorononanoic acid (6:3-FTCA, CAS 27854-30-4, purity ≥ 97%), 2,H,2H,3H,3H-Perfluoroundecanoate (8:3-FTCA, CAS 83310-58-1, purity ≥ 97%), 2H,2H,3H,3H-Perfluorohexansulfonic acid (4:2-FTSA, CAS 757124-72-4, purity ≥ 97%) and 2H,2H,3H,3H-Perfluorooctane-1-sulfonate (6:2 FTSA, CAS 59587-39-2, purity ≥ 97%) were from Synquest Laboratories (Alachua, FL). HSA (CAS 70024-90-7, purity ≥ 95%, fraction V fatty acid free) and hexadecanoic acid (CAS 57-10-3, natural, purity ≥ 98%) were from Millipore Sigma (Burlington, MA). HEPES (4-(2-hydroxyethyl)-1-piperazineethanesulfonic acid), sodium chloride, methanol, dimethylsulfoxide, decanoic acid (CAS 334-48-5, purity ≥ 99%) and ibuprofen (CAS 15687-27-1, purity ≥ 99%), and potassium chloride (KCl, CAS 7447-40-7, purity ≥ 99.7%) were purchased from Thermo Fisher (Waltham, MA).

**Figure 1.**
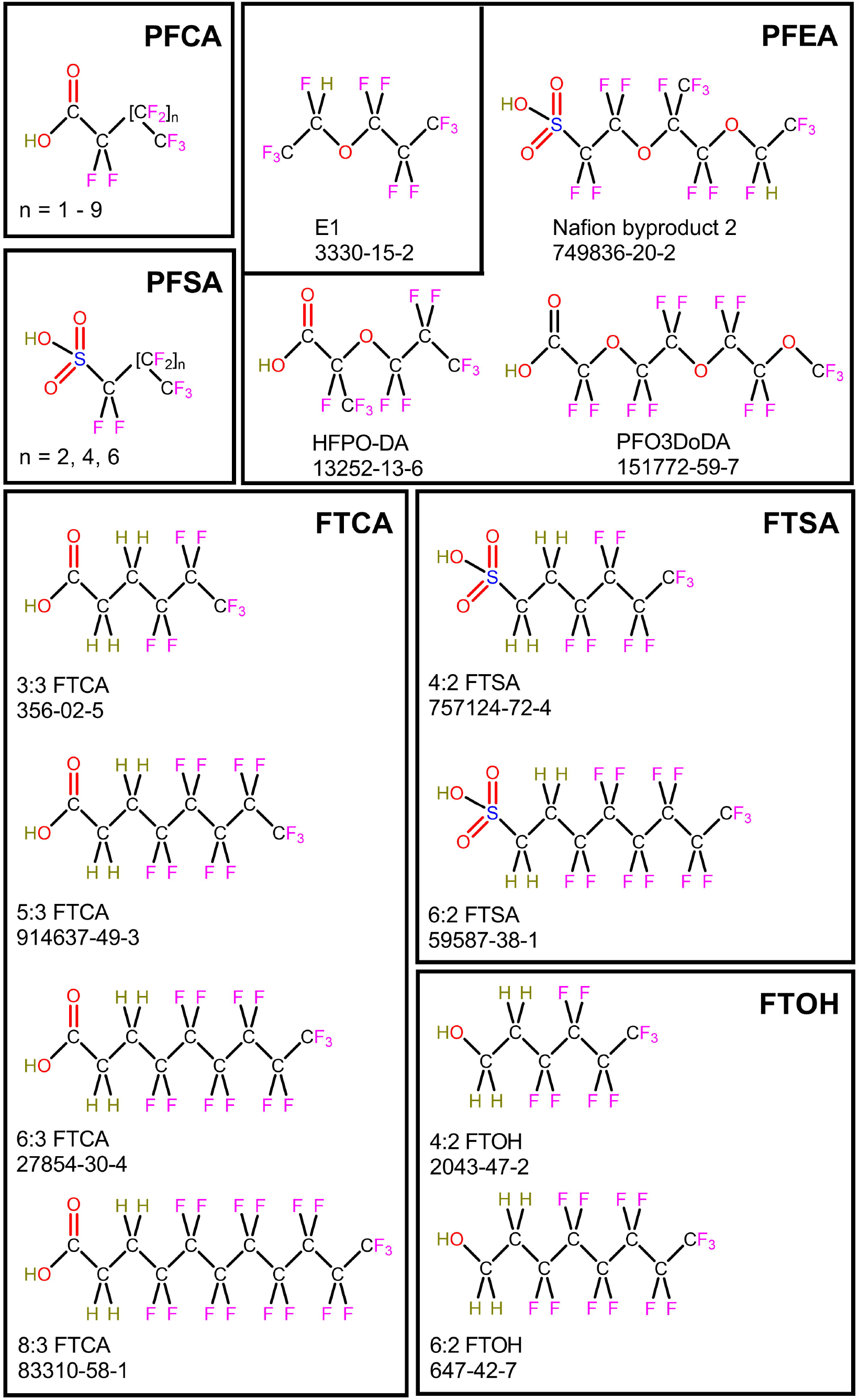
Structures of PFAS analyzed.

### 2.2 Control and Test Chemical Preparation

Stock solutions (20 mM) of PFBA, PFPeA, PFHxA, PFHpA, PFOA, PFBS, PFHxS, PFOS, HFPO-DA, Nafion bp2, 6:3-FTCA, 6:2-FTSA, decanoic acid, ibuprofen, and KCl were prepared in aqueous 1x HEPES buffered saline (HBS, 140 mM NaCl, 50 mM HEPES, 0.38 mM Na_2_HPO_4_, pH 7.2). A 1:1 mixture of HBS and DMSO was used as a solvent for PFNA, PFDA, PFunDA, and 8:3-FTCA stocks due to limited aqueous solubility, and the fatty acids and warfarin were dissolved into HBS supplemented with 30% methanol. For experiments evaluating possible solvent effects 20 mM stock solutions of PFOA were prepared in all three solvents. The HBS concentrations used in solvents containing DMSO or methanol were adjusted to ensure that the final concentration of the thermal denaturation buffer contained 140 mM NaCl, 50 mM HEPES, 0.38 mM Na_2_HPO_4_. Solution pH for PFAS stocks were confirmed to be 7.4 and stocks were stored at -20° C. For thermal stability concentration response analysis, stock solutions were serially diluted into solvent. Stocks of HSA (1 mM) were prepared in 2x HBS and then diluted with an equal volume of H_2_O to final desired concentrations.

### 2.3 Differential Scanning Fluorimetry

Temperature control and fluorescence detection were performed using a Step One Plus Real-Time PCR System (Applied Biosystems; Grand Island, NY) with indicator dye (GloMelt) fluorescence (*λ*_Ex_ = 468 nm *λ*_Em_ = 507 nm) detected using the FAM/SYBR filter set and the passive reference dye carboxyrhodamine (*λ*_Ex_ = 588 nm *λ*_Em_ = 608 nm) detected using the ROX channel. Thermal denaturation was performed in sealed optical 96-well reaction plates (MicroAmp Fast, Applied Biosystems) using the following conditions: 10 minutes at 37° C for one holding stage, followed by a ramp profile from 37° C to 99° C at a rate of 0.2° C/sec.

Following optimization, each DSF assay contained 0.125 mM HSA in a final volume of 20 μl. Stock solutions of each test chemical were serially diluted into HBS, with final concentrations ranging from 50 µM to 10 mM. Working fluorophore solutions (200x in 0.1% DMSO) diluted 1:20, and ROX (40 μM) diluted 1:10 were prepared immediately prior to each experiment with 2 μl of each used for each assay. At least two independent plates were run for each experimental unit. Controls run on each plate included matching vehicle control (no ligand; KCl added for potassium salts), no protein control, and a minimum of three concentrations of decanoic acid as a positive control for protein stabilization. To evaluate the sensitivity of the assay to detect DMSO mediated conversion of HPFO-DA to E1^22^, HFPO-DA was prepared in a 1:1 mixture of HBS and DMSO and maintained at room temperature for 4 hours before experimental analysis. To evaluate whether volatile compounds were entering the gas phase to reduce concentrations of PFAS, experiments were performed using different reaction volumes ranging from 10 µL to 200 µL in each well for 4:2-FTOH, 6:2-FTOH, PFHxS, and 6:2-FTS.

### 2.4 Data analysis and statistics

All presented DSF data are representative of multiple experiments each containing 3 replicates for each sample. Matching vehicle blank controls lacking test compound were included on the same plate for each experiment. Raw thermocycler data were exported to Excel (Microsoft) and statistical analysis was performed using SPSS v26 (IBM, Armonk, NY) or GraphPad Prism (v8.3.0, GraphPad Software Inc., San Diego, CA). Data are reported as mean values ± SD following background subtraction. Assay data is reported in relative fluorescent light units (RFU). The T_m_ is defined as the temperature at which the maximum change in fluorescence is observed, indicating half of the protein is unfolded. PFAS concentration response curves were smoothed using the Savitzky and Golay method ^23^, EC_50_ estimates are derived using a 4-parameter variable slope model, and dissociation constants were calculated using a single site ligand binding model using the formula ^24^:

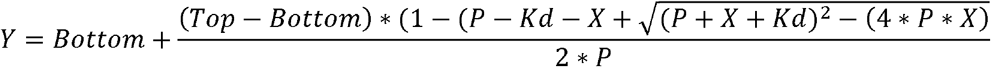

Top is the maximal response, bottom is minimal response, P is protein concentration, K_d_ is dissociation constant, X is ligand concentration, and Y is change in T_m_. This equation requires that a maximal response be detected, which is limited by the solubility of the compounds of interest. This equation fits a concentration-response curve to the melt shift and provides an estimated dissociation constant. Using this equation, the calculated K_d_ is most accurate when its value is greater than 50% of the protein concentration and requires ligand concentrations approximately ten times the K_d_ ^24^.

The relationship between number of aliphatic carbons or number of fluorine and the binding affinity of HSA for each compound was determined using a second order polynomial (quadratic) best fit with least squares regression. Comparison between protein concentrations and comparisons of calculated binding affinities between different compounds was performed using one-way analysis of variance (ANOVA) and a Tukey’s post hoc test was performed to evaluate pair-wise differences. Significance between differences in values was defined as *p* < .05.

## 3. Results

### 3.1 Thermal melt assay optimization

Concentrations of HSA between 0.05 mM to 0.625 mM were evaluated to identify the HSA concentration that yielded maximal signal to noise ratio (Figure 2A). The observed T_m_ for HSA (71.3^°;^C) did not vary across the concentration range analyzed (F (4, 10) = 2.19, p = .14; Figure 2B). Optimal performance was for assays containing 0.125 mM HSA (Figure 2A). Including an initial 10-minute preincubation at 37^°;^ C decreased the relatively high initial fluorescence observed for HSA, and the optimal temperature ramp rate was determined to be 0.2° C/sec. Most study compounds were sufficiently soluble to use 1x HBS as a solvent for 20 mM stock solutions. The limited aqueous solubility of the C9-C11 PFCA and 8:3-FTCA required use of HBS containing 50% DMSO, and the fatty acids and warfarin required using 30% methanol as a solvent. Possible solvent effects were investigated for PFOA that was solubilized in each of the three solvents. Assay results for HSA binding of PFOA binding were not significantly influenced by the stock solution solvent (F (2, 15) = 0.005, p = .996) (Table 1). The increase in potassium ions from the potassium salts of PFHxS, PFOS, 8:3-FTCA, and 6:2-FTSA did not affect assay results (data not shown).

**Figure 2:**
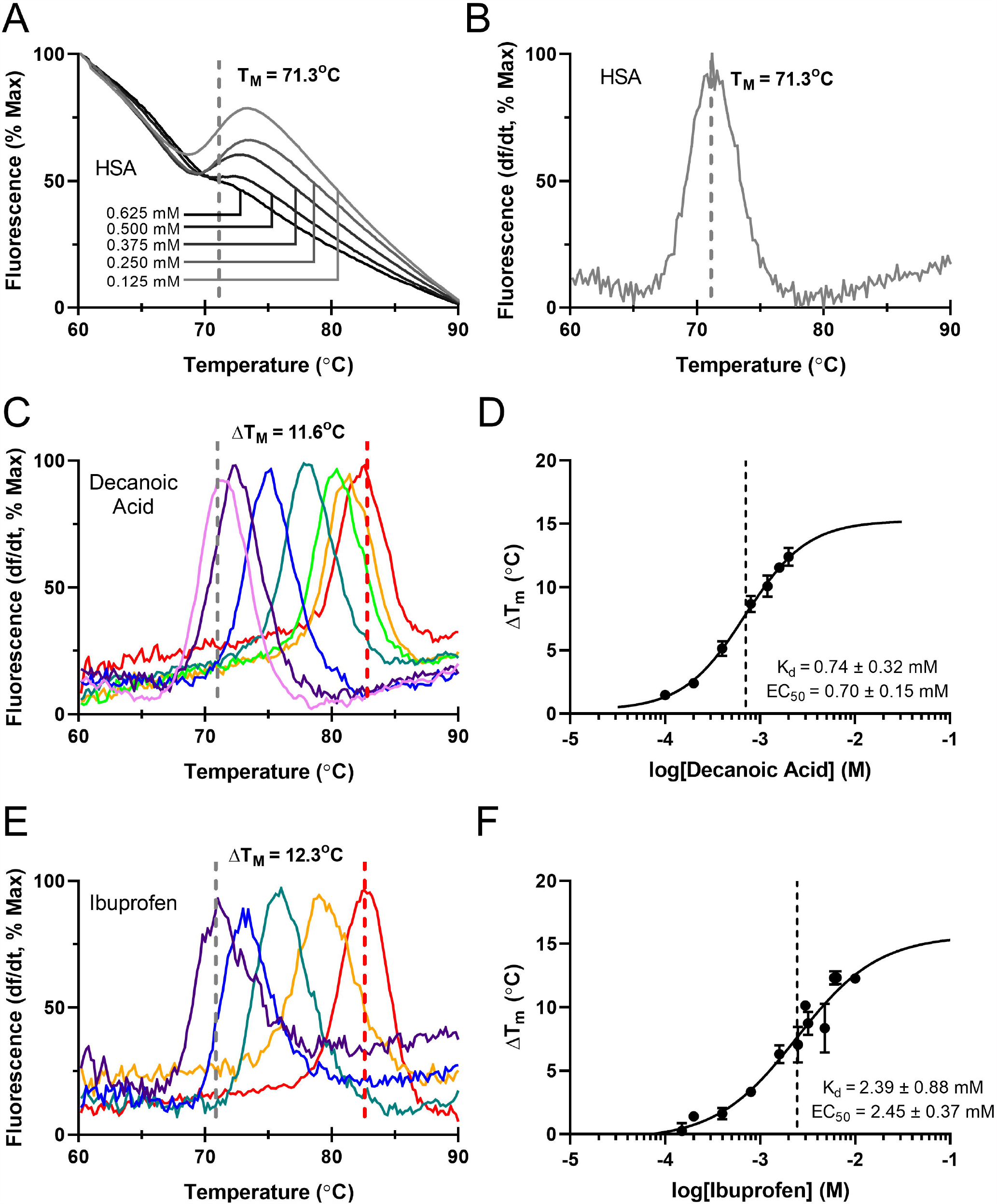
Validation of DSF for measuring control compound binding. The fluorescence of HSA alone, normalized to the % maximum, as temperature was increased from 60-90°C is shown with the melting temperature indicated as the point at which half of the protein is inferred to be unfolded (A). Increasing concentrations of HSA (0.125 mM to 0.625 mM) from light gray to black are shown. The fluorescence signal of concentrations below 0.125 mM was not detectable. The derivative fluorescence of HSA alone, plotted as the derivative of fluorescence divided by the derivative of time, as temperature was increased from 60-90°C is shown with the melting temperature indicated as the maximum of the derivative curve (B). Derivative fluorescent curves for HSA with the fatty acid decanoic acid (C) or known albumin binding compound ibuprofen (E) as temperature was increased from 60-90°C, are shown with increasing concentrations of compound indicated by increasing wavelength of color from violet to red. The maximum change in temperature for HSA alone is shown between the dashed gray and red lines. The regression of the change in temperature plotted against the logarithmic transformed concentration, in molar units, is shown for decanoic acid (D) and ibuprofen (F), with the log(EC_50_) indicated by a dashed line. n ≥ 3 across at least two replicate plates for all compounds.

**Table 1.** Analysis of solvent effects.

| Solvent | $K_d$ (mM) | EC50 (mM) | $\Delta T_m$ (°C) |
| --- | --- | --- | --- |
| HEPES-buffered saline (HBS) | $0.83 \pm 0.27$ | $0.84 \pm 0.10$ | $13.5 \pm 0.26$ |
| Methanol (30% in HBS) | $0.83 \pm 0.15$ | $0.78 \pm 0.17$ | $13.5 \pm 0.19$ |
| DMSO (50% in HBS) | $0.84 \pm 0.20$ | $0.85 \pm 0.02$ | $13.2 \pm 0.35$ |

$\Delta T_m$ is °C, EC50 and $K_d$ are mean values reported in ± SD. Each compound was run on at least two separate plates with $n \geq 4$

### 3.2 Measurement of HSA binding affinity for known HSA binding compounds

Octanoic acid, decanoic acid, hexadecenoic acid, warfarin, and ibuprofen were used as positive controls to evaluate whether DSF estimates of binding affinities were comparable to published values using other methods. Analysis of the fatty acid-induced melting temperature shift of HSA determined a K_d_ of 2.10 ± 0.47 mM for octanoic acid, 0.74 ± 0.32 mM for decanoic (Figure 2C and 2D), and 0.030 ± 0.02 for hexadecanoic acid (Table 2). Two-way ANOVA revealed that the fatty acids were significantly different (F (2, 15) = 63, p < .0001), with Tukey’s post-hoc comparison indicating that each fatty acid was significantly different from the other two examined. The calculated K_d_ for HSA binding of ibuprofen was 2.39 ± 0.88 mM (Figure 2E and 2F) and warfarin was 0.16 ± .10 mM (Table 2). The calculated affinities of HSA binding for each of all compounds are within the range of previously determined values.^32-35^

**Table 2.** Binding affinity of HSA for control compounds.

| Compounds | CAS ID | $R^2$ | $\Delta T_m$ (°C) | EC50 (mM) | $K_d$ (mM) |
| --- | --- | --- | --- | --- | --- |
| Octanoic Acid | 124-07-2 | 0.93 | $3.1 \pm 0.17$ | $2.15 \pm 0.25$ | $2.10 \pm 0.47$ |
| Decanoic Acid | 334-48-5 | 0.98 | $12.4 \pm 0.50$ | $0.70 \pm 0.15$ | $0.74 \pm 0.32$ |
| Hexadecanoic Acid | 57-10-3 | 0.95 | $7.2 \pm 0.13$ | $0.084 \pm 0.05$ | $0.03 \pm 0.02$ |
| Ibuprofen | 15687-27-1 | 0.97 | $12.3 \pm 0.24$ | $2.45 \pm 0.37$ | $2.39 \pm 0.88$ |
| Warfarin | 81-81-2 | 0.97 | $9.3 \pm 0.21$ | $0.19 \pm 0.06$ | $0.16 \pm 0.10$ |

$\Delta T_m$ is °C, EC50 and $K_d$ are mean values reported in ± SD. Each compound was run on at least two separate plates with $n \geq 4$

### 3.3 Measurement of HSA binding affinity for PFAS

Numerous studies have evaluated albumin binding of PFOA and PFOS.^26–31^ Using DSF, the calculated K_d_ for HSA binding of PFOA was 0.83 ± 0.38 mM (Figure 3A and 3B), and 0.69 ± 0.078 mM for PFOS (Figure 3C and 3D; Table 3). The calculated K_d_ for HSA binding of PFOA and PFOS were similar to previously reported values, although these values vary greatly depending on the method and assay conditions.^26–31^ The findings from the DSF assay and calculated dissociation constant for each PFCA (C4-C12), PFSA (C4-C8), the ether-containing PFAS, (PFAE; Figure 3E and 3F), and eight fluorotelomer compounds are shown in Table 3. It is notable that the fluorotelomer alcohols 4:2 FTOH and 6:2 FTOH were not bound by HSA and that fluorotelomer compounds with a carboxylate or sulfonate charged group were bound by HSA at affinities similar to those observed for PFAA with the same number of aliphatic carbons (Table 3).

**Figure 3:**
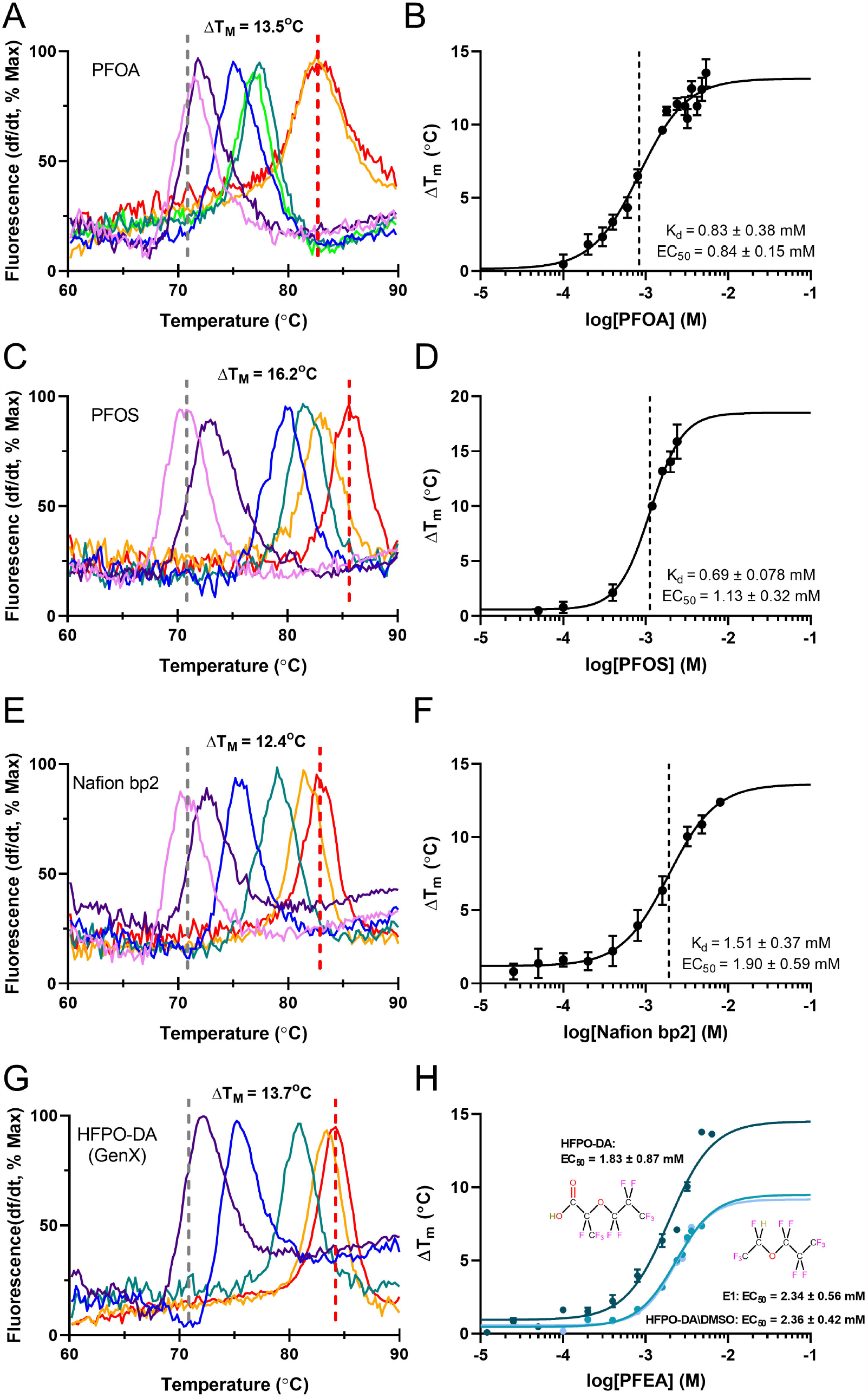
Validation of DSF for measuring PFAS binding. Derivative fluorescent curves for HSA with the PFAA PFOA (A), PFOS (C), and Nafion byproduct 2 (E), HFPO-DA (GenX) (G), as temperature was increased from 60-90°C, are shown with increasing concentrations of compound indicated by increasing wavelength of color from violet to red. The maximum change in temperature from HSA alone is shown between the dashed gray and red lines. The regression of the change in temperature plotted against the logarithmic transformed concentration, in molar units, is shown for PFOA (B), PFOS (D), Nafion byproduct 2 (F), and GenX (H) with the log(EC_50_) indicated by a dashed line. n ≥ 3 across at least two replicate plates for all compounds. In panel (H), the regression of the change in temperature plotted against the logarithmic transformed concentration, in molar units, is also shown for GenX in DMSO and E1, along with chemical structures for GenX and E1.

**Table 3.** Binding affinitv of HSA for PFAS.

| PFCA | Cas ID | Chain Length | Aliphatic Carbons | Fluorines | $R^2$ | $\Delta T_m$ (°C) | EC50 (mM) | Kd (mM) |
| --- | --- | --- | --- | --- | --- | --- | --- | --- |
| PFBA | 375-22-4 | 3 | 4 | 7 | 0.94 | $6.48 \pm 0.18$ | $2.61 \pm 0.47$ | $2.57 \pm 0.78$ |
| PFPeA | 2706-90-3 | 4 | 5 | 9 | 0.97 | $13.1 \pm 0.066$ | $2.14 \pm 0.42$ | $2.10 \pm 0.56$ |
| PFHxA | 307-24-4 | 5 | 6 | 11 | 0.98 | $10.6 \pm 0.24$ | $1.40 \pm 0.27$ | $1.64 \pm 0.47$ |
| PFHpA | 375-85-9 | 6 | 7 | 13 | 0.95 | $15.3 \pm 0.41$ | $0.68 \pm 0.15$ | $0.44 \pm 0.20$ |
| PFOA | 335-67-1 | 7 | 8 | 15 | 0.97 | $13.5 \pm 0.41$ | $0.84 \pm 0.15$ | $0.83 \pm 0.38$ |
| PFNA | 375-95-1 | 8 | 9 | 17 | 0.97 | $13.4 \pm 0.34$ | $0.60 \pm 0.23$ | $0.58 \pm 0.21$ |
| PFDA | 335-76-2 | 9 | 10 | 19 | 0.99 | $17.2 \pm 0.13$ | $1.11 \pm 0.17$ | $1.19 \pm 0.59$ |
| PFUnDA | 2058-94-8 | 10 | 11 | 21 | 0.98 | $9.02 \pm 0.47$ | $1.49 \pm 0.048$ | $1.36 \pm 1.06$ |
| PFDoA | 307-55-1 | 11 | 12 | 23 | 0.98 | $7.74 \pm 0.20$ | $2.51 \pm 0.34$ | $1.89 \pm 0.64$ |
| **PFSA** |  |  |  |  |  |  |  |  |
| PFBS | 375-73-5 | 4 | 4 | 9 | 0.96 | $11.9 \pm 0.11$ | $1.72 \pm 0.56$ | $1.65 \pm 0.69$ |
| PFHxS | 3871-99-6 | 6 | 6 | 13 | 0.98 | $11.0 \pm 0.24$ | $0.98 \pm 0.069$ | $0.71 \pm 0.47$ |
| PFOS | 2795-39-3 | 8 | 8 | 17 | 0.98 | $16.2 \pm 0.90$ | $1.13 \pm 0.32$ | $0.69 \pm .078$ |
| **Per- and Polyfluorinated Alkyl Ethers** |  |  |  |  |  |  |  |  |
| E1 | 3330-15-2 | 5 | 5 | 11 | 0.93 | $7.32 \pm 0.059$ | $2.34 \pm 0.56$ | $1.64 \pm 0.34$ |
| HFPO-DA | 13252-13-6 | 5 | 5 | 11 | 0.97 | $13.7 \pm 0.17$ | $1.83 \pm 0.87$ | $1.60 \pm 0.55$ |
| Nafion bp2 | 749836-20-2 | 7 | 7 | 14 | 0.98 | $12.4 \pm 0.17$ | $1.90 \pm 0.59$ | $1.51 \pm 0.37$ |
| PFO3DoDA | 151772-59-7 | 6 | 7 | 13 | 0.95 | $20.6 \pm 0.82$ | $1.67 \pm 0.47$ | $1.53 \pm 0.34$ |
| **Fluorotelomer Alcohols** |  |  |  |  |  |  |  |  |
| 4:2 FTOH | 2043-47-2 | 4 | 6 | 9 | N/A | $0 \pm 0$ | N/A | N/A |
| 6:2 FTOH | 647-42-7 | 6 | 8 | 13 | N/A | $0 \pm 0$ | N/A | N/A |
| **Fluorotelomer Carboxylic Acids** |  |  |  |  |  |  |  |  |
| 3:3 FTCA | 356-02-5 | 3 | 6 | 7 | 0.81 | $2.24 \pm 0.062$ | $2.06 \pm 0.51$ | $1.71 \pm 0.47$ |
| 5:3 FTCA | 914637-49-3 | 5 | 8 | 11 | 0.94 | $3.62 \pm 0.14$ | $1.48 \pm 0.24$ | $0.62 \pm 0.10$ |
| 6:3 FTCA | 27854-30-4 | 6 | 9 | 13 | 0.95 | $9.50 \pm 0.20$ | $0.84 \pm 0.21$ | $0.81 \pm 0.23$ |
| 8:3 FTCA | 83310-58-1 | 8 | 11 | 17 | 0.89 | $10.01 \pm 0.17$ | $1.16 \pm 0.42$ | $0.97 \pm 0.17$ |
| **Fluorotelomer Sulfonic Acids** |  |  |  |  |  |  |  |  |
| 4:2 FTSA | 757124-72-4 | 4 | 6 | 9 | 0.93 | $3.49 \pm 0.12$ | $1.45 \pm 0.34$ | $1.07 \pm 0.47$ |
| 6:2 FTSA | 59587-38-1 | 6 | 8 | 13 | 0.91 | $3.60 \pm 0.47$ | $0.47 \pm 0.29$ | $0.37 \pm 0.34$ |

Chain length is the number of per/poly fluorinated carbons; $\Delta T_m$ is °C, EC50 and Kd are mean values reported in ± SD. Each compound was run on at least two separate plates with $n \geq 4$

To determine whether the high volatility of the fluorotelomer alcohols was responsible for the absence of albumin binding, values were determined for 4:2-FTOH, 6:2-FTOH, PFHxS, and 6:2-FTS at volumes of 10, 20, 50, and 200 µL that resulted in different volumes of gaseous phase in each sealed reaction well. At 200 µL, the well is with no gas phase. There were no differences in the thermal shift profile at different volumes for any of the four PFAS measured, findings that suggest volatility of the fluorotelomer alcohols was not responsible for the lack of albumin binding (4:2-FTOH, F(3, 8) = 0.90, p = .48; 6:2-FTOH, F(3, 8) = 0.14, p = .93; PFHxS, F(3, 8) = 0.63, p = .61; 6:2-FTSA, F(3, 8) = 0.67, p = .60).

To investigate the sensitivity of the assay to distinguish binding properties for closely related compounds, we compared assay results for HFPO-DA prepared in aqueous buffer or in DMSO containing buffer with assay results for E1 directly. In DMSO, HFPO-DA is rapidly converted to E1 via decarboxylation.^22^ Two-way ANOVA of the area under the curve of the concentration-response curves for HFPO-DA in DMSO, HFPO-DA in buffer alone (Figure 3G), and E1 reveals significant differences (F (2, 29) = 144, p < .0001), with Tukey’s post-hoc analysis indicating that HFPO-DA in DMSO is indistinguishable from the E1 curve with EC_50_ values of 2.34 ± 0.56 mM and 2.36 ± 0.42 mM, respectively (p = .98; Figure 3H).Tukey’s post-hoc analysis found that HFPO-DA in buffer alone is significantly different from HFPO-DA in DMSO and E1 in buffer (both p < .0001).

### 3.4 Physiochemical determinants of HSA binding

To interrogate in more detail determinants of HSA binding of PFAS, the relationship between calculated binding affinities, and the number of per- and polyfluorinated carbons, number of aliphatic carbons, or total fluorine numbers for the PFCA series from C4-C12 and across all compounds were analyzed. Except for the PFAE compounds, highest affinity was observed for compounds containing 6-8 fluorinated carbons, 7-9 aliphatic carbons, and containing 13-17 fluorine (Figure 4). For the PFAE, a simple linear regression was more appropriate. For the PFCA series from C4-C12, the best-fit curve for binding affinity by number of per- and polyfluorinated carbons was = 6.30 - 1.50*X* + 0.10*X*^2^ (Figure 4A; R^2^ = 0.88) and across all compounds except PFAE was = 4.73 - 1.08*X* + 0.074*X*^2^ (R^2^ = 0.54). For PFAE, the simple linear regression by per- and polyfluorinated carbons was = -0.02X + 1.7 (Figure 4B; R^2^ = 0.79). Except for the PFAE, the best-fit curve for the number of aliphatic carbons was = 6.52 - 1.39*X* + 0.083*X*^2^ (Figure 4C; R^2^ = 0.69) and by number of fluorine was = 5.35 - 0.58*X* + 0.019*X*^2^ (Figure 4D; R^2^ = 0.54). For the PFAE family, the linear regression by number of aliphatic carbons was = -0.06X+1.9 (Figure 4C; R^2^ = 0.52) and by number of fluorine was = -0.01X+1.7 (Figure 4D; R^2^ = 0.77).

**Figure 4:**
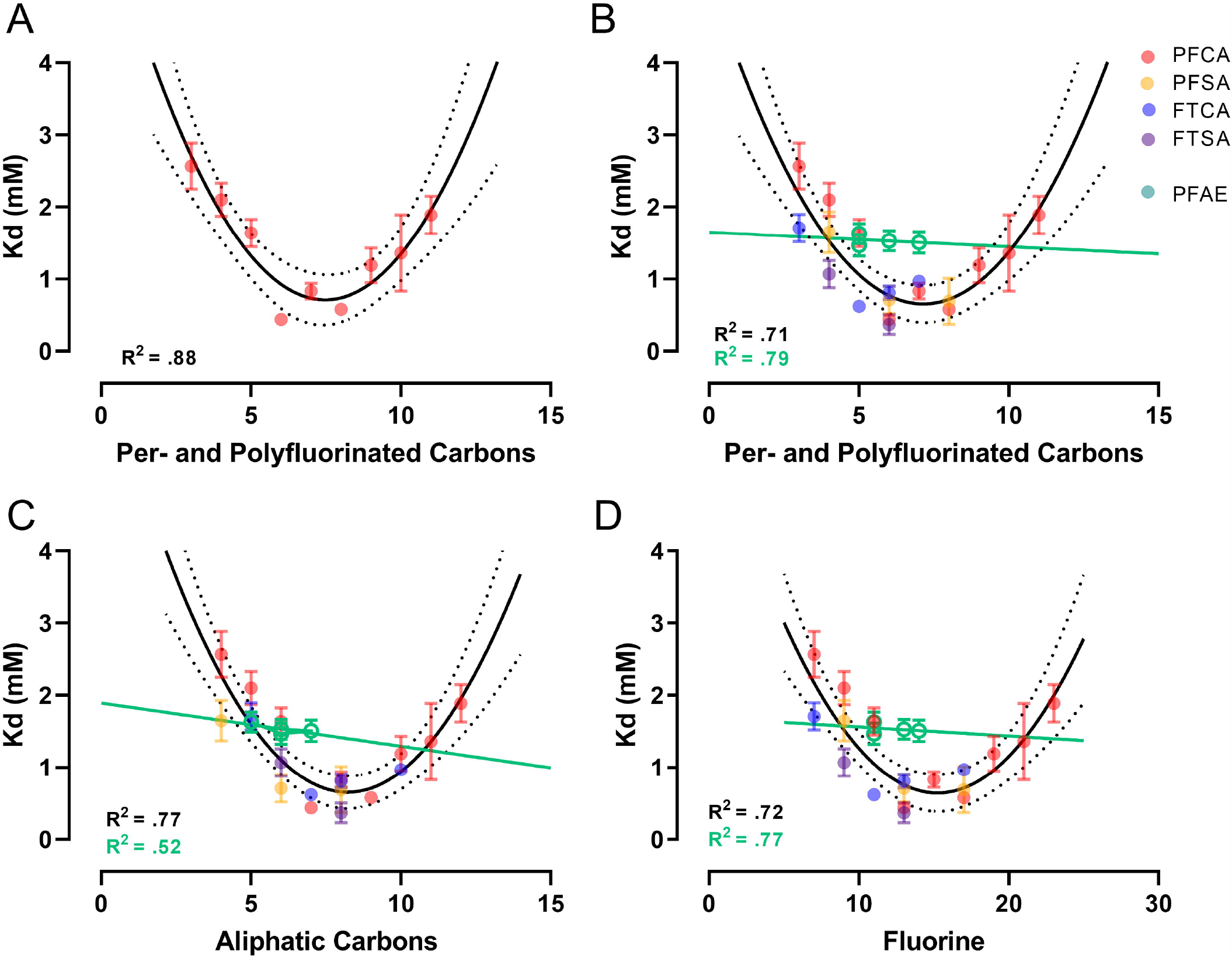
Effect of carbon chain length and fluorine moieties on PFAS binding. The binding affinity of the PFCA (A) and all analyzed PFAA and PFAE (B, C, and D) are plotted against the number of per- and polyfluorinated carbons (A, B), aliphatic carbons, (C), or fluorine (D). For all PFAA except PFAE, a quadratic line of best fit with 95% confidence interval in dashed lines was generated using least squares regression. Each class is indicated by different colors, with PFCA in red, PFSA in orange, PFAE in green, FTCA in blue, and FTSA in purple. N ≥ 3 across at least two replicate plates for all compounds.

## 4. Discussion

### 4.1 Optimization and demonstration of assay utility

The goal of the current studies was to develop a rapid, high-throughput assay capable of measuring protein binding affinity of a diverse collection of PFAS compounds. The presented experiments describe the optimization and use of a DSF assay for assessing HSA binding properties for control compounds known to bind albumin and 24 PFAS from six subclasses. Critical initial experiments aimed to optimize DSF for measuring PFAS binding included determination of optimal protein and dye concentrations to maximize signal to noise ratio. Those efforts were found especially critical for determining albumin binding due to its multiple surface accessible hydrophobic binding sites that increased baseline fluorescence.^32^ Additional key factors analyzed during assay development included use of a HEPES buffer to ensure that PFAS with low pKa did not affect assay pH, maintaining consistent ionic strength, determination of appropriate solvents, and optimization of assay temperature ramp rates. Results of those initial experiments identified appropriate conditions for determining the binding affinities of structurally diverse sets of natural fatty acids, small molecule pharmaceuticals, and multiple subclasses of PFAS in a rapid (less than 3 hour) format. The accuracy and reproducibility of the binding affinities calculated using DSF was demonstrated for known albumin-binding drugs warfarin and ibuprofen, C10-C16 fatty acids, PFOA and PFOS.^25–27,29,31,33-36^ Further demonstrating the utility of this DSF thermal shift approach, comparative evaluation of the HSA binding affinities of structurally diverse subclasses of PFAS revealed that functional groups, number of aliphatic carbons, and number of fluorine bonded to carbons were among the key physiochemical properties that influenced binding.

### 4.2 Impacts of physiochemical properties on HSA binding affinity

Published K_d_ values for HSA binding of fatty acids, drugs, and PFAS are variable and can span many orders of magnitude.^25–27,29,31,33,35-38^ Because the absolute K_d_ values depend on the specific experimental conditions of each assay, it is most useful to compare relative affinities across different assays. The pattern of HSA affinity for fatty acids observed here is consistent with previous findings that found affinity increased with longer chain length such that the affinity of hexadecanoate > decanoate > octanoate.^33,37,38^ For these fatty acids, increasing chain length allows the methylene tails to extend further into the deep hydrophobic cavities of HSA, with HSA binding sites completely filled by fatty acids of length C18-C20.^39^ While HSA can bind fatty acids longer than C20, binding affinity is decreased because the methylene tails are not fully accommodated and therefore have lower binding energies than optimal C16-C20 fatty acids.^39^ Some PFAS, specifically PFCA, have structural similarities with fatty acids, and the high-affinity fatty acid binding sites are likely sites for PFAS interactions.^40^ Because PFCAs are fatty acids with fluorine replacing the aliphatic hydrogens, the same properties that allow albumin to bind fatty acids also allow albumin to bind PFAS. However, unlike fatty acids, PFAS have fluorinated alkyl tails that impart oleophobic amphiphilic surfactant properties and decrease the relative water solubility of PFAS ^41^. Because of these complexities, numerous physiochemical properties, including the number of per- and polyfluorinated carbons, the number of aliphatic carbons, the number of fluorine attached to aliphatic carbons, and the functional headgroups were evaluated for their influence on relative binding affinities of HSA for PFAS. Within each class of analyzed PFAS, HSA relative affinity for aliphatic carbon length was: C4-C5 < C6-C9 > C10+.The optimal structure for binding with HSA appears to be between six and nine aliphatic carbons. Unlike fatty acids, the increasing aliphatic backbone of C10+ PFAS appears to prevent optimal binding due to an increase in net negative charge resulting in oleophobic steric hindrance that may force the longer-chain PFAA to fold.^40^ Consistent with these observations, molecular docking experiments predict that PFAA with more than 11 carbons cannot easily fit into the binding pocket of fatty acid binding protein, but these molecular docking studies became less reliable for predicting HSA affinity for PFCA >9 perfluorinated carbons due to a lack of experimental affinity data.^42^ Ng and Hungerbuehler specifically emphasize the critical need for further experimental data on which to base molecular docking simulations, and the assay described here can provide this data via rapid comparison of protein affinity for multiple compounds assayed using the same experimental conditions.^42^

The importance of the functional headgroup in the affinity of HSA for PFAS was evaluated by comparing binding affinity between fluorotelomer compounds with an alcohol headgroup to those with a carboxylate or sulfonate headgroup. Strikingly, the two fluorotelomer alcohols tested, 4:2-FTOH and 6:2-FTOH, did not bind HSA. The fluorotelomer compounds with a carboxylate or sulfonate group were bound by HSA with affinities comparable to PFAA, demonstrating that the charged functional group is important for HSA binding. Those findings are consistent with complexation energy analysis demonstrating the fluorinated chain of PFOA and PFOS interacted significantly with the aliphatic portion of the positively charged guanidinium groups of Arg 218 and Arg 222 and the backbone amine group of Asn 294, and these interactions were essential in the overall complexation between HSA and PFAS.^40^ However, it is important to note that E1, an ether PFAS with no charged functional group, was also bound by albumin. It is likely that E1, and potentially other PFAE, are bound by albumin via a different mode than the other PFAS. This hypothesis is consistent with the binding patterns of fluorinated ether anesthetics, where there is evidence of nonpolar binding in subdomain IIIB by enflurane, a fluorinated ether anesthetic with a nominal dipole that contrasts with the polar binding by similar compounds with larger dipole moments (e.g. isoflurane).^43^

When comparing compounds with the same number of per- and polyfluorinated carbons but different functional groups, the pattern of binding affinity followed the pattern: ether acids < carboxylic acids < sulfonic acids. This pattern applied when comparing PFCA to PFSA and FTCA to fluorotelomer sulfonic acids. Previous reports demonstrate that the longer perfluorinated chain of PFOS provides greater complexation energy than PFOA, whereby apolar interactions account for much more of the binding between HSA and PFOS via increased van der Waals interactions.^40^ This observation appears to hold true across classes, and increased van der Waals interactions provided by the additional fluorinated carbon in the PFSA of equal chain length to the PFCA are explain the increased affinity of HSA for sulfonated moieties.

Similarly, HSA had higher affinity for the fluorotelomer acids than the PFAA with equal numbers of per- and polyfluorinated carbons, providing further evidence that number of aliphatic carbons is providing increased stability with HSA by increasing the fit into the hydrophobic binding pockets. Finally, the findings that albumin had lower affinity for the PFEA than PFAA with the same number of per- and polyfluorinated carbons are consistent with previous work demonstrating that linear PFAS bind albumin much more strongly than their branched isomers, potentially reflecting that ether linkages impart structures similar to those adopted by branched isomers.^31^

### 4.3 Strengths and limitations

The DSF method utilized here has numerous advantages over typical methods including titration chemistry or surface plasmon resonance; namely, DSF requires substantially less protein (0.08 mg of HSA per assay) and the assay can be completed and provide affinity data for up to 8 PFAS compounds in less than four hours using the 96 well format. Ongoing studies have demonstrated that the assay is scalable to a 384 well format to further increase throughput. Additionally, DSF is performed using real-time PCR instruments that are widely available and accessible by most laboratories ^44^. Further, this assay can be easily adapted to analyze binding affinities for a wide array of purified proteins and assay conditions ^16,45,46^. It is important to note that DSF assays often employ the hydrophobic fluorophore SYPRO Orange, SYPRO Orange is not compatible with assays containing detergents or surfactants and is not useful for analyzing PFAS due to the amphipathic surfactant properties of many PFAS. The assay described here was optimized to use an alternative environment sensing fluorophore because of anticipated limitations of SYPRO Orange, namely the surfactant and detergent-like properties of many PFAS would render the hydrophobic dyes incompatible.^24^ Preliminary analysis found that a number of commercially available fluorescent rotor dyes, including dicyanovinyl)julolidine, 9-(2-Carboxy-2-cyanovinyl)julolidine, 4-(4-(dimethylamino)styryl)-N-methylpyridinium iodide, and the used dye preparation GloMelt(tm) were compatible for DSF analysis of PFAS (not shown).

An additional strength of this DSF assay is its ability to detect changes in PFAS chemistry, evidenced by the ability to detect the conversion of HFPO-DA to E1 following incubation in DMSO. Previous analysis has demonstrated that use of DMSO as a solvent for HFPO-DA results in rapid and complete conversion of HFPO-DA to E1 in under four hours.^22^ Using this DSF assay, the complete decarboxylation of HFPO-DA by DMSO was demonstrated by the observed differences in the concentration response relationship differences between HFPO-DA in HEPES-buffered saline and HFPO-DA in DMSO. The concentration response curve and the resulting EC_50_ and HSA binding affinity values for HFPO-DA in DMSO were found identical to that of E1 demonstrating the quantitative decarboxylation of HFPO-DA to E1.

Whereas we have demonstrated that PFAS compounds in aqueous solutions or prepared in the solvent methanol or DMSO were compatible with this assay, the limited aqueous solubility of C12 and longer PFCA and other longer chain PFAS did not allow analysis across the concentration range needed to accurately determine binding affinities for HSA. Because the complete range of concentration-response must be determined to accurately evaluate the binding affinities and associated parameters, the DSF assay is limited to PFAS with sufficient solubility in aqueous solutions. Additionally, binding affinities determined using the DSF method are generated over a range of temperatures and are not directly related to dissociation constant values determined using other methods^47^. The ΔT_m_ used to calculate K_d_ has the advantage of giving a more complete view of the thermodynamic system when comparing compound binding. Consistent with previous reports that binding affinities calculated using DSF are often lower than using other methods due to calculating the affinity at melting temperature instead of physiological temperature, the absolute affinities of HSA for PFAS were lower but within the same order of magnitude of published values ^24^. The differences in reported values are at least partly due to the fact that the dissociation constant is determined at the higher melting temperature of the protein with ligand, rather than at a constant temperature of 20° or 37° C typically used for other methods.^34^

With these results, we have shown the utility of a rapid and sensitive high throughput DSF assay that is able to define protein-binding affinities and identify physiochemical properties involved in protein binding for large numbers of PFAS. This proof-of-concept study was focused on the major serum transport protein albumin because of its critical role in PFAS distribution and bioaccumulation. However, because of the flexibility of this assay, PFAS binding properties of other purified proteins from any species of interest can be evaluated. Key parameters identified as determinants of PFAS HSA binding of included the constitutive functional groups and the number of aliphatic carbons. Disruption of the aliphatic chain was found to decrease HSA binding affinity and potentially alter the modes of binding. This was especially evident for the tetrafluoroethyl ether E1, which lacked a charged functional group but unlike fluorotelomer alcohols, was bound by HSA, finding that suggest binding of this short chain PFAS may be similar to HSA binding of volatile fluoroether anesthetics. Adaptation of the DSF methods demonstrated here will allow rapid characterization of protein affinity for PFAS, improve computational modeling of protein-PFAS binding kinetics, and allow prioritization of PFAS for subsequent toxicity evaluation.

---

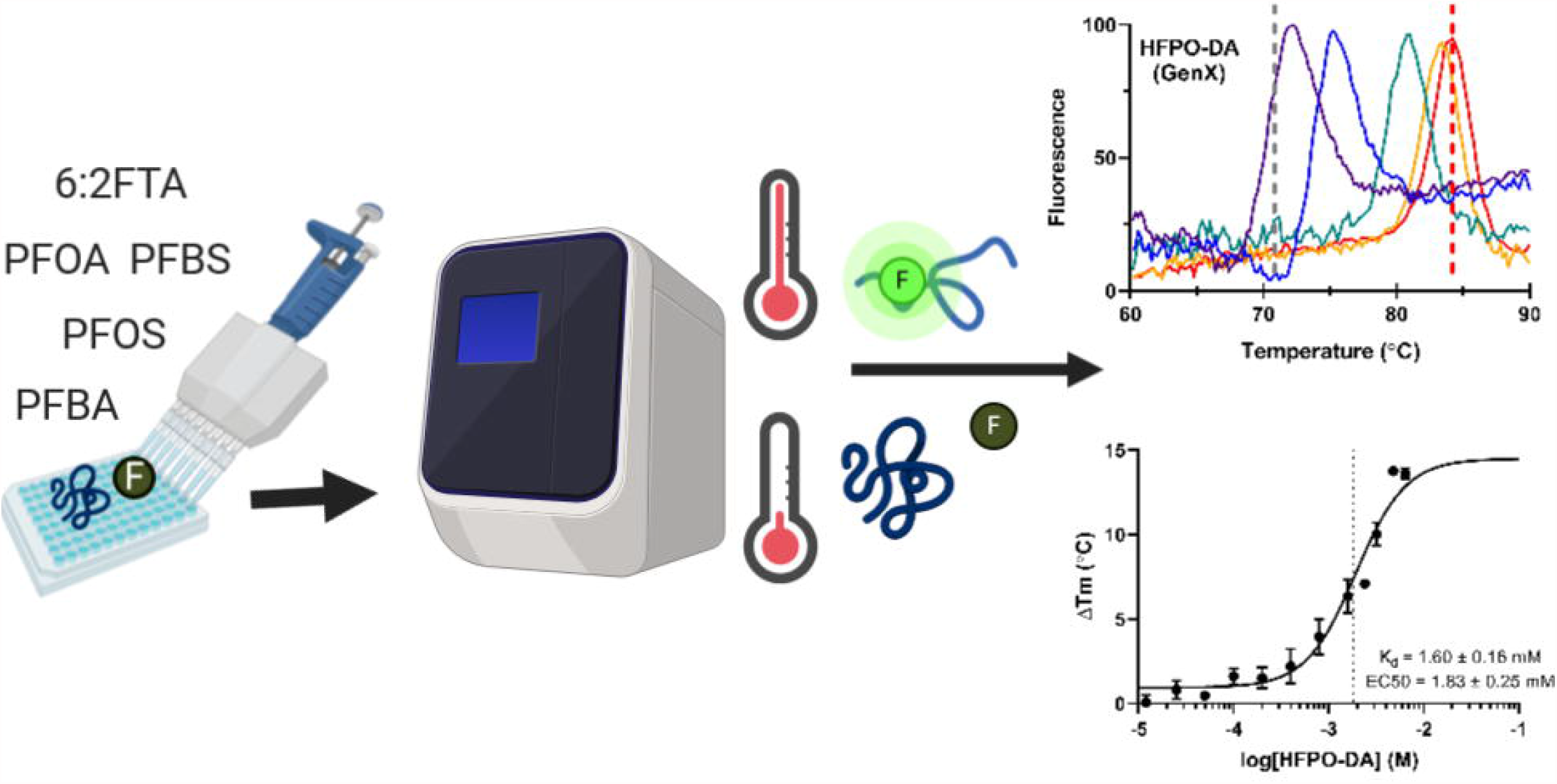

